# The effects of thermal alteration on organic matter bioavailability in deeply buried marine sediments

**DOI:** 10.64898/2026.06.16.732717

**Authors:** Samuel M. McNichol, Sunita R. Shah Walter, Andreas Teske, Nagissa Mahmoudi

**Author notes:** **Corresponding Author:** Nagissa Mahmoudi.

## Abstract

A substantial fraction of marine sediments experience elevated temperatures due to burial or hydrothermal activity. These conditions can fundamentally reshape both microbial activity and the chemical nature of sedimentary organic matter (OM). Laboratory incubations have demonstrated that moderate heating of marine sediments can lead to the production of labile organic compounds such as acetate, however, it remains unclear whether heating alters the bioavailability of the remaining OM pool. In this study, we experimentally tested the effect of temperature on the bioavailability of OM through a series of bioreactor experiments using deeply buried sediment collected from Guaymas Basin (Gulf of California). We measured acetate concentrations in sterilized Guaymas Basin sediments before and after artificial heating (70°C for 7 days) to quantify abiotic acetate generation. We then conducted incubations of a model marine bacterium with sterilized, artificially heated sediment and tracked respired CO_2_ production and its associated ^13^C and ^14^C signatures. Our study revealed that sediment depth and hydrothermal history strongly control abiotic acetate production, with higher acetate yields from shallower, cooler sediments. Respiration rates in control and heated sediment incubations were nearly identical, indicating that heating does not measurably alter the bioavailability of bulk sedimentary OM. Moreover, the δ^13^C values of respired CO_2_ were indistinguishable between control and heated sediment incubations while the Δ^14^C values were more depleted in the first 24 hours in incubations with heated sediment. This transient offset suggests that low-temperature heating mobilizes a small pool of older material due to desorption of mineral-bound OM without altering overall bioavailability. Our findings shed light on the role of thermal alteration in shaping carbon cycling in marine sediments by influencing how OM is made available to sedimentary microorganisms.

## 1. Introduction

Marine sediments cover more than two thirds of the Earth’s surface and store vast amounts of organic matter (OM) that is continually transformed by diverse microbial communities. The transformation and remineralization of OM after its deposition on the seafloor arises from tightly coupled interactions between microorganisms, geochemical conditions, and physical processes. While much of our understanding of these processes comes from low-temperature environments, a substantial portion of marine sediments experience elevated temperatures as a result of burial or hydrothermal activity. An estimated 35% of global marine sediment exists at temperatures of 60°C or higher (LaRowe et al., 2017). Such conditions can fundamentally reshape both microbial activity and the chemical nature of OM, thereby influencing microbial pathways and rates of carbon cycling in marine sediments.

Thermal alteration can begin at temperatures as low as ~50°C (Tissot and Welte, 1984) and proceeds initially through disruption of weak bonds that bind organic molecules onto mineral surfaces. With increasing temperatures, these processes promote the generation and mobilization of low molecular-weight (MW) compounds, including organic acids such as acetate (Lang et al., 2019). Where there is advective flow, these compounds can be transported upward through the sediment column once produced, supporting microbial life in overlying sediments (Amend and Teske, 2005; Zhuang et al., 2019; Brünjes et al., 2025). For example, sediments collected near buried hydrothermal sills have a much lower percent total organic carbon (TOC) than organic rich surface sediment above (Lizarralde et al., 2023), reflecting extensive thermal alteration and mobilization of OM. In parallel, OM is continuously transformed by microbial processing following deposition and throughout burial. Microorganisms consume substrates from the bulk OM pool while also producing organic compounds such as organic acids (e.g., acetate) via fermentative pathways (Burdige, 2006). Together, these abiotic and biotic processes interact to shape the balance between OM degradation and preservation in sediments.

Previous laboratory studies have demonstrated that artificial heating of sediments can generate small organic compounds in both freshly deposited and deeply buried sediments. Early incubation experiments using shallow estuarine and deep subsurface sediments heated to temperatures of up to 100°C observed rapid release of acetate into porewaters over timescales ranging from days (Wellsbury et al., 1997) to years (Parkes et al., 2007). Subsequent work using high-resolution mass spectrometry demonstrated that prolonged heating (~90 °C) of shallow marine sediments generates not only organic acids, but a chemically diverse pool of dissolved organic compounds, including amino acids (Lin et al., 2017). More recent incubations of subsurface sediments across temperatures ranging from 20°C to 85°C suggest that the combined effects of microbial and hydrothermal processing can partition the sedimentary OM pool. Some of the OM becomes more bioavailable via the production of low-MW organic compounds, while a second pool becomes more condensed and less accessible to microorganisms (Gan et al., 2025). While these studies collectively have revealed that heating generates small organic compounds, the impact of heating on the bioavailability of the remaining sedimentary OM pool remains unclear. This distinction is critical because these labile, low-MW compounds are vulnerable to rapid mobilization upward through the sediment column (Lin et al., 2017), rather than being retained and consumed at the depths at which they are produced (Knoke et al., 2025). Consequently, microbial communities residing in deeper sediments must rely on the remaining, thermally altered OM pool. Thus, it remains unresolved whether heating merely generates labile compounds that subsequently are mobilized or whether it also alters the bioavailability of the remaining OM pool that supports microbial populations at depth.

Here, we tested how elevated temperatures influence the bioavailability of sedimentary OM by conducting a series of bioreactor experiments in which a marine bacterial isolate was incubated with sterilized sediment that had been artificially heated in the laboratory followed by removal of volatile compounds. Specifically, we measured microbially respired CO_2_ and its associated ^13^C and ^14^C signatures to track OM pools supporting microbial respiration. Experiments were conducted using sediments from shallow and deep burial depths collected from Guaymas Basin, located in the Gulf of California. Guaymas Basin sediments experience hydrothermal heating which enabled us to evaluate how burial depth and sediment heating history modulate the effects of heating on OM bioavailability. In parallel, acetate concentrations were measured in sterilized sediments before and after heating as a proxy for the suite of low MW organic compounds generated during heating. This allowed us to assess the extent to which abiotic acetate production is governed by the freshness of OM, which declines with both sediment age and past exposure to hydrothermal heating. Our findings shed light on the role of temperature in shaping carbon cycling in marine sediments by influencing how OM is transformed and made available to microbes.

## 2. Materials & Methods

### 2.1 Study site and sampling

Guaymas Basin is a young, active spreading center in the central Gulf of California where sediments are impacted by hydrothermal heating (Figure 1). Heating in Guaymas Basin occurs in both on-axis and off-axis regions when magma intrudes laterally into the sediment column creating steep thermal gradients (up to 1000 °C/km). Exposure of sediments to elevated temperatures leads to the production of petroleum hydrocarbons and organic acids (Simoneit, 1985; Lin et al., 2017; Zhuang et al., 2019). Due to highly productive overlying waters and riverine inputs of terrestrial OM, Guaymas Basin has a thick sediment column characterized by high organic carbon content (typically 3-4% TOC) (Teske et al., 2016).

**Figure 1.**
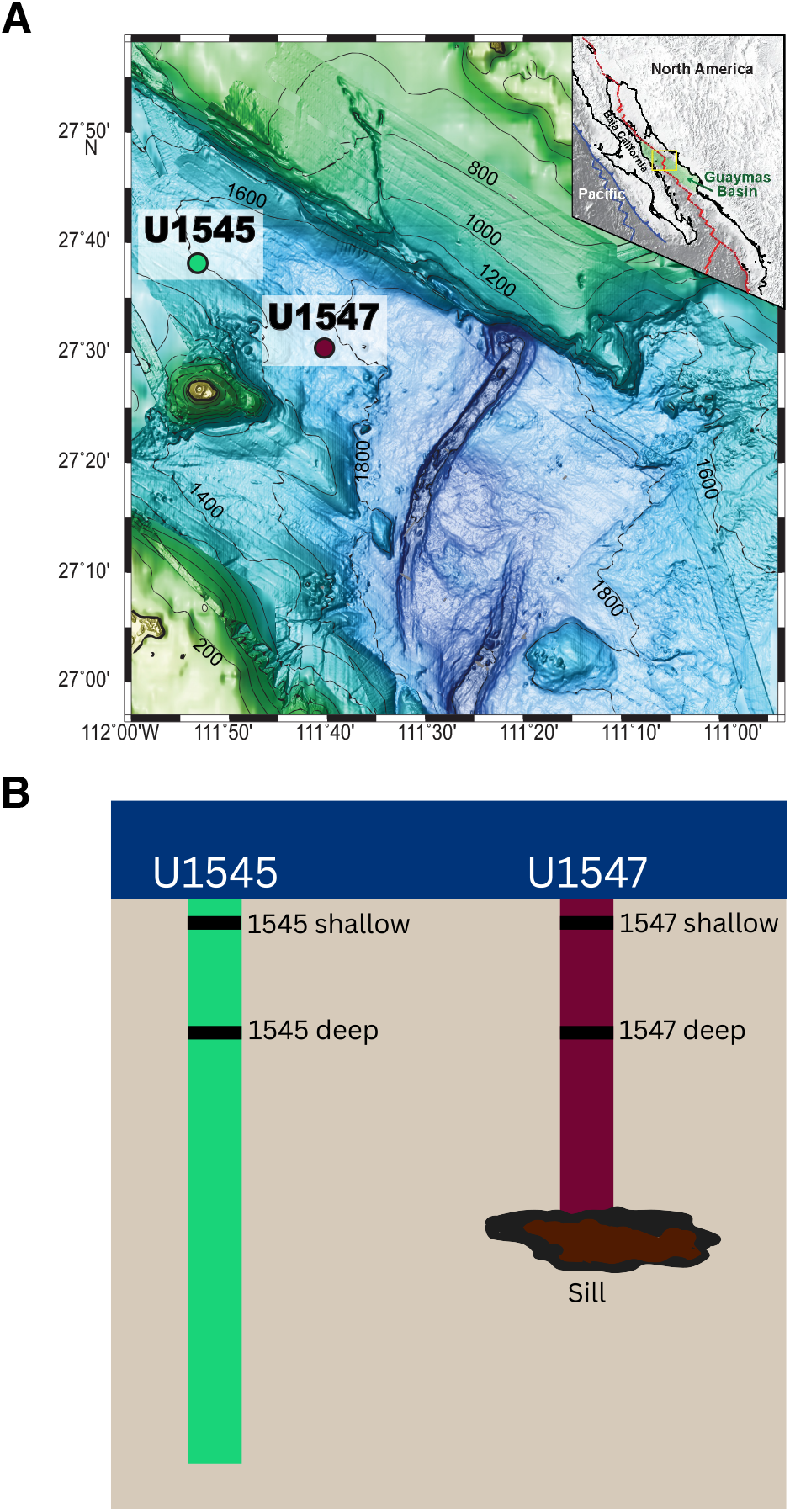
Bathymetry of Guaymas Basin (Gulf of California), with (A) sites U1545 and U1547 used in this study indicated by green and red circles (Adapted from Teske et al., 2021). (B) Illustration of the locations of sites U1545 and U1547 relative to sills emplaced within the sediment column in Guaymas Basin (Adapted from Bojanova et al., 2023).

Sediment samples selected for this study were obtained during IODP Expedition 385 (September 15 – November 15, 2025) from drill core sites U1545 and U1547 (Figure 1). Cores were frozen at −80°C immediately after collection. Details on sample processing, collection, and in situ measurements are outlined in the IODP 385 Proceedings (Teske et al., 2021). Heated, off-axis sediment was collected from drill site U1547, where sediments are exposed to a temperature gradient of 500-1000°C/km due to a shallow, recently emplaced hot sill at ~115 meters below seafloor (mbsf) (Teske et al., 2019, 2021). Both shallow (8 mbsf) and deep (56 mbsf) core sections were selected, with *in situ* temperatures of 13°C and 41°C (Table S1). Off-axis sediment from site U1545 was collected as a regional background site, as there are no magmatic sills directly beneath the drill core location. Site U1545 however still experiences a temperature gradient of 200-250°C/km, the background gradient for much of Guaymas Basin (Teske et al., 2021). Shallow (4 mbsf) and deep (57 mbsf) core sections were selected, with temperatures of 7°C and 17°C, respectively (Table S1).

### 2.2 Sediment heating and sterilization

Core sections were homogenized and titrated to pH 2-2.5 with 10% HCl in an ice bath to remove carbonate species. Subsequently, homogenized sediments were freeze dried and sterilized by gamma-irradiation via a ^137^Cs source (radioactive cesium) to receive a total dose of ~40 kGy. A fraction of the freeze dried, sterilized sediment sample was heated in the laboratory. Prior to heating, sediments were saturated with modified Tibbles-Rawlings (T-R) minimal medium containing no carbon source (Tibbles and Rawlings, 1994). The slurry was then transferred to a combusted glass jar, sealed with Parafilm, capped and heated in a drying oven (F Air 2.3CF, VWR) at 70°C for seven days. Subsequently, the jar of sediment was frozen at −20°C and incubated in the bioreactor within ~24 hours of completing the heating.

### 2.3 Acetate concentrations before and after heating

Acetate concentrations were quantified in sediment slurries before (control) and after heating, to quantify abiotic generation of acetate in sediments collected from different depths and hydrothermal histories. A total of 3 ml of sediment slurry was filtered (PES filter, 0.22 µm, VWR) in replicate and immediately frozen at −20°C before and after heating. Acetate concentrations were determined using gas chromatography with a flame ionization detector (GC-FID) in the Resnick Water and Environment Lab at the California Institute of Technology. Filtered sediment slurry samples were thawed to room temperature and 500 µl of sample, 100 µl of 10% HCl, and 100 µl of ethanol were added to a 10 ml headspace vial (Thermo Fisher) and tightly sealed. The sealed samples were incubated at 95°C for 3 h in a sand bath to derivatize ethyl acetate. After incubation, the headspace was analyzed by gas chromatography (GC) with a flame ionization detector (FID) (Agilent 6890) using a HP-Plot/Q column (30 m, 0.32 mm, 20 µm) and a headspace injector (HP 7964). Acetate concentrations were quantified in samples based on a calibration curve created from sodium acetate. Ethyl acetate was derivatized from sodium acetate standard solutions with each batch of samples heated in the sand bath.

### 2.4 IsoCaRB incubation experiments

Control and heated sediment samples were incubated in the Isotopic Carbon Respirometer Bioreactor (IsoCaRB) system to quantify microbial respiration and the isotopic (δ^13^C and Δ^14^C) composition of respired CO_2_. This system is comprised of a gas delivery and purification system, a custom 3 l Pyrex culture vessel, an inline CO_2_ detector and integrated LabVIEW data-logging program, custom CO_2_ traps, and a vacuum extraction line. All incubations were conducted using established procedure for the IsoCaRB system, including sterilization and assembly, and CO_2_ collection and purification (Beaupre et al., 2016). CO_2_ concentrations obtained during incubations are corrected for the confounding effects of mixing in the culture vessel headspace and decreasing media volume to constrain the rate of CO_2_ generation per unit volume of growth medium (ug C L^−1^ min^−1^), which serves as a proxy for the microbial CO_2_ production rate (Mahmoudi et al., 2017). The culture vessel is also equipped with a sampling port that allows for direct sampling of the culture medium throughout incubation.

Parallel incubations of control and heated sediments were conducted at room temperature in the IsoCaRB system with a model marine bacterium, *Pseudoalteromonas* sp. 3D05 (Figure S1). This bacterial isolate was originally obtained from coastal ocean water samples (Datta et al., 2016) and is capable of degrading a wide range of complex substrates (McNichol et al., 2024; Baumser et al., 2024). Its genome encodes for extracellular hydrolytic enzymes needed to break down both proteins and carbohydrates (Mahmoudi et al., 2020). In addition, *Pseudoalteromonas* sp. 3D05 encodes for ATP-binding cassette (ABC) transporters and Ton-B dependent transporters, which facilitate the uptake of amino acids, peptides, sugars and carbohydrates directly into the cell (Schauer et al., 2008; How et al., 2025). Cells were prepared for inoculation into the bioreactor as published previously (Mahmoudi et al., 2020). Briefly, cells were grown from frozen glycerol stocks and cell density was monitored by measuring optical density (OD) at 600 nm, based on a calibration curve between OD and colony forming units (CFUs). Once cells reached a desired cell density of ~ 2.5 × 10^7^ CFU/ml, 50 ml of cell culture was harvested by centrifugation for 10 min at 3000 xg and washed two times with T-R minimal medium containing no carbon source. The resulting cell pellet was injected into the IsoCaRB system using a 3 ml syringe and a 20-gauge needle. Cell density was measured frequently (every ~12 hours) during the incubation by manual counting of colony forming units (CFUs) on agar plates from subsamples of the sediment slurry.

Equal masses of sediment were used in control and heated sediment incubations to ensure that the mass of organic carbon was consistent across experiments. Prior to inoculation, the bioreactor was sparged with CO_2_-free helium for 48-96 hours. This served two important purposes for the experiments. First, it purged the system of residual atmospheric CO_2_ and ensured that all measured CO_2_ reflects microbial respiration, and that the isotope signals are not obscured by atmospheric signatures. Second, sparging removed most of the volatile, low MW organic compounds that are generated during heating. Removal of acetate was confirmed through measurement of the initial acetate concentrations in the bioreactor slurry (Table S2). This allowed us to directly test the impact of heating on the bioavailability of the remaining sedimentary OM pool. Lastly, all incubations were conducted under oxic conditions (20% O_2_) to ensure that microbial respiration was not limited by electron acceptors.

### 2.5 Isotopic analysis of microbially respired CO_2_

Microbially respired CO_2_ was continuously collected during IsoCaRB incubations as successive fractions in custom molecular sieve traps. The duration of each CO_2_ fraction ranged from 7-26 h to ensure that no fraction contained more than 2 mg of carbon. The collected CO_2_ fractions were sent to the National Ocean Sciences Accelerator Mass Spectrometry (NOSAMS) Facility at the Woods Hole Oceanographic Institution for Δ^14^C analysis. An aliquot of each sample was split for ^13^C measurement at the Oregon State Stable Isotope Collaboratory (OSSIC) Facility at the University of Oregon. The remainder of the CO_2_ was reduced to graphite for ^14^C measurement by accelerator mass spectrometry (AMS) (Vogel et al., 1987). All isotopic data are corrected for background contamination associated with the IsoCaRB system as previously described (Mahmoudi et al., 2020). Stable isotope values (^13^C) are reported versus the VPDB standard. Radiocarbon values (^14^C) are reported in Δ^14^C notation, where Δ^14^C is the relative deviation from the ^14^C/^12^C ratio of the atmosphere in 1950 corrected for kinetic fractionation using measured δ^13^C values (Stuiver and Polach, 1977).

## 3. Results

### 3.1 Abiotic acetate generation from heating

Across all sediment samples, short-term heating (70°C for 7 days) led to the generation of acetate. Both sediment depth and hydrothermal history were important factors in the amount of acetate production. For both cores, sediments from shallower depths had higher acetate yields. With site U1545 sediments, heating produced 3.9 ± 0.6 mg acetate per g organic carbon (OC) in shallow sediment and 3.0 ± 0.2 mg acetate per g OC in deep sediment (Figure 2). The effect of depth was more pronounced in site U1547 sediments, where heating generated 4.0 ± 0.9 mg acetate per g OC in shallow sediment but only 1.1 ± 0.3 mg acetate per g OC in deep sediment (Figure 2). This indicates that, as OM is exposed to elevated temperatures during long-term burial, acetate generating precursors are removed from the OM pool. Moreover, it affirms that abiotic acetate production is governed by the freshness of OM, which declines with both sediment age and past exposure to hydrothermal heating.

**Figure 2.**
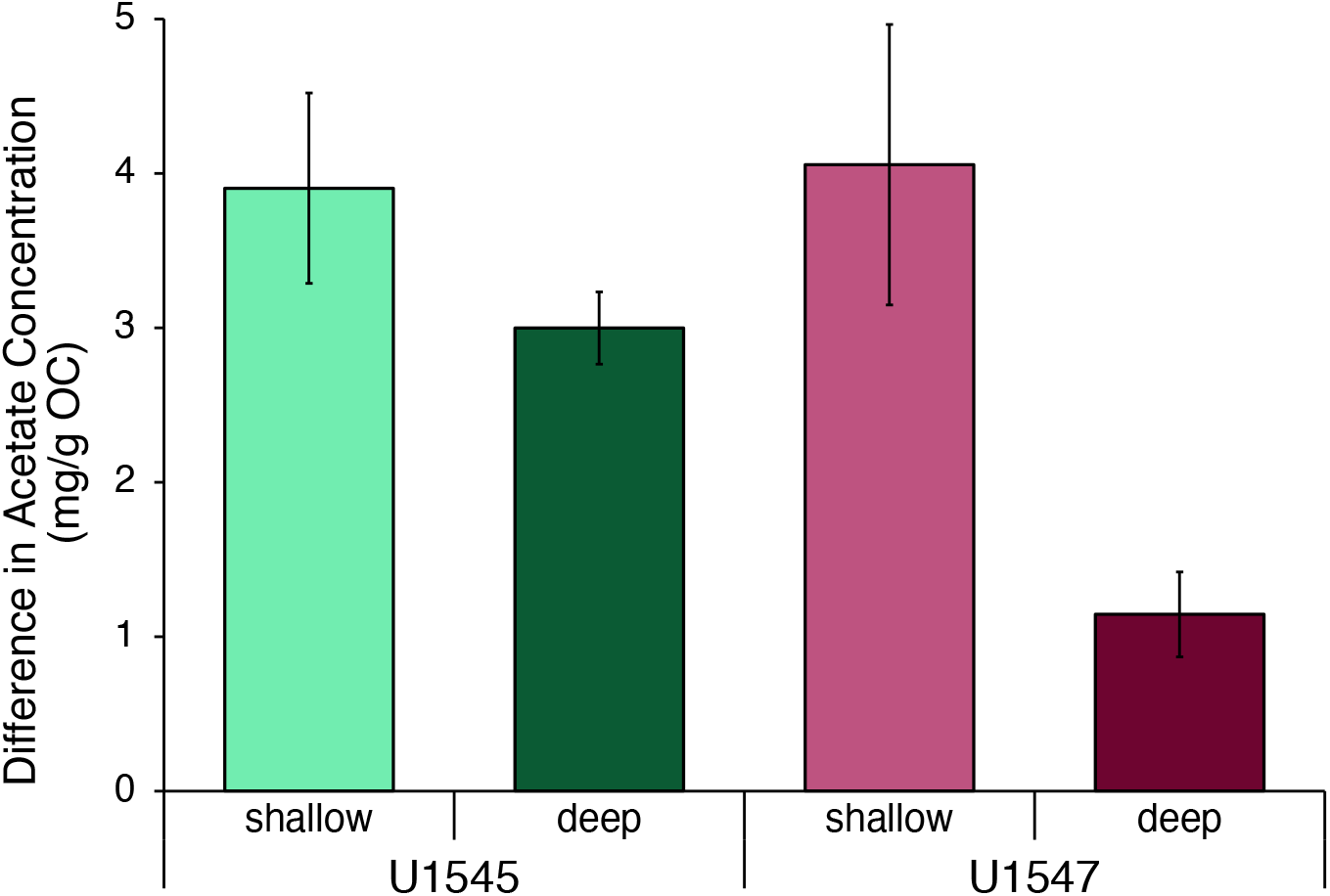
(A) Difference in acetate concentration (mg/g OC) in sediment slurries before and after short-term heating (70°C, 7 days). Acetate concentration was determined by GC-FID from filtered subsamples of sediment slurry.

### 3.2 Microbial remineralization of sedimentary OM before and after heating

Total CO_2_ yield, respiration rates, and cell density were quantified during bioreactor incubations with control and heated sediments. To facilitate comparisons in total C respired between sites and depths, total respired C was normalized to the mass of organic carbon (OC) in each incubation (Table S3) to account for differences in sediment mass across incubations. The normalized total yield of CO_2_ respired was nearly identical between incubations with heated and control sediments, a pattern that was consistent across all sediment depths and sites (Figure 3). In shallow sediments from site U1545, 1.3% OC was respired in both control and heated sediment incubations, while deep U1545 sediments showed similarly small differences; 1.1% OC respired in the control vs. 1.2% OC respired after heating. Site U1547 sediments exhibited the same trend, such that incubation of shallow sediment yielded 0.55% and 0.64% OC respired in control and heated sediment incubations, respectively, and deep sediments resulted in 0.73% OC and 0.76% OC respired (Figure 3).

**Figure 3.**
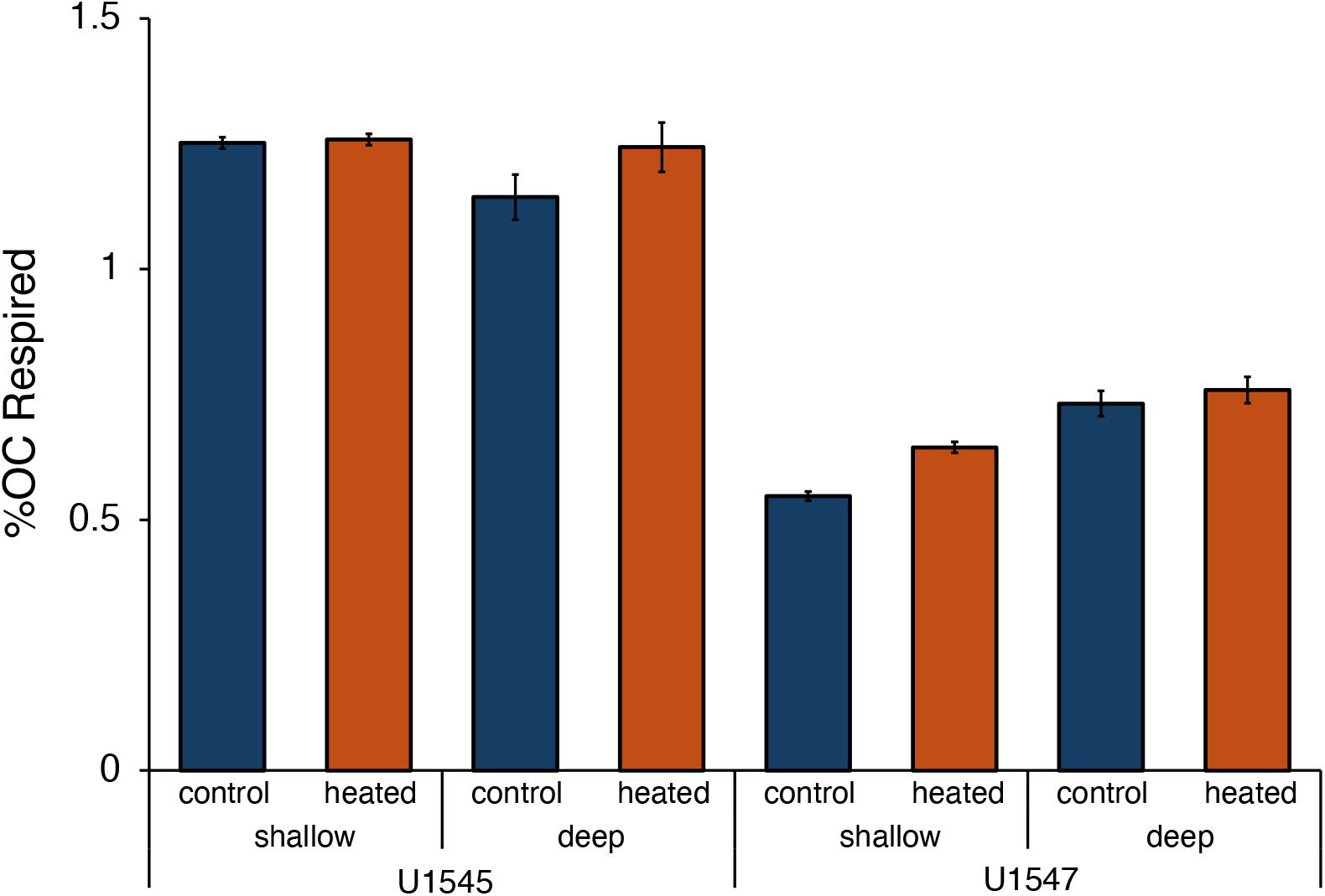
Percent OC respired in bioreactor incubations with control and heated sediments; total C respired was normalized to the mass of OC in each incubation.

Consistent with normalized CO_2_ yields, the respiration dynamics and cell growth patterns were similar in control and heated sediment incubations. Across all incubations, CO_2_ production rates increased rapidly with the onset of incubation, peaking within the first 12 h before gradually decreasing to near-baseline values within ~6 days (Figures 4, S2). Maximum respiration rates were comparable between control and heated sediment incubations; peak respiration rates reached 1.1 µg C L^−1^ min^−1^ and 0.9 µg C L^−1^ min^−1^ with shallow and deep sediments from site U1545 respectively, irrespective of heating. In incubations with shallow sediments from site U1547, respiration rates reached 1.5 ug C L^−1^ min^−1^ and 1.9 ug C L^−1^ min^−1^ with control and heated sediments, and 1.2 ug C L^−1^ min^−1^ with deep sediment, independent of heating (Figures 4, S2). Across all incubations, cell abundances increased over the first 12 to 27 h, peaking between 6 to 7 × 10^7^ CFU ml^−1^ (Figures S3, S4; Table S4). Following this increase, cell density either remained steady or began slowly declining after the first 24 h. By ~4 days, cell density across all incubations had fallen below initial inoculation levels in all incubations. Collectively, these results indicate that short-term heating does not substantially alter the bioavailability of the remaining OM pool.

**Figure 4.**
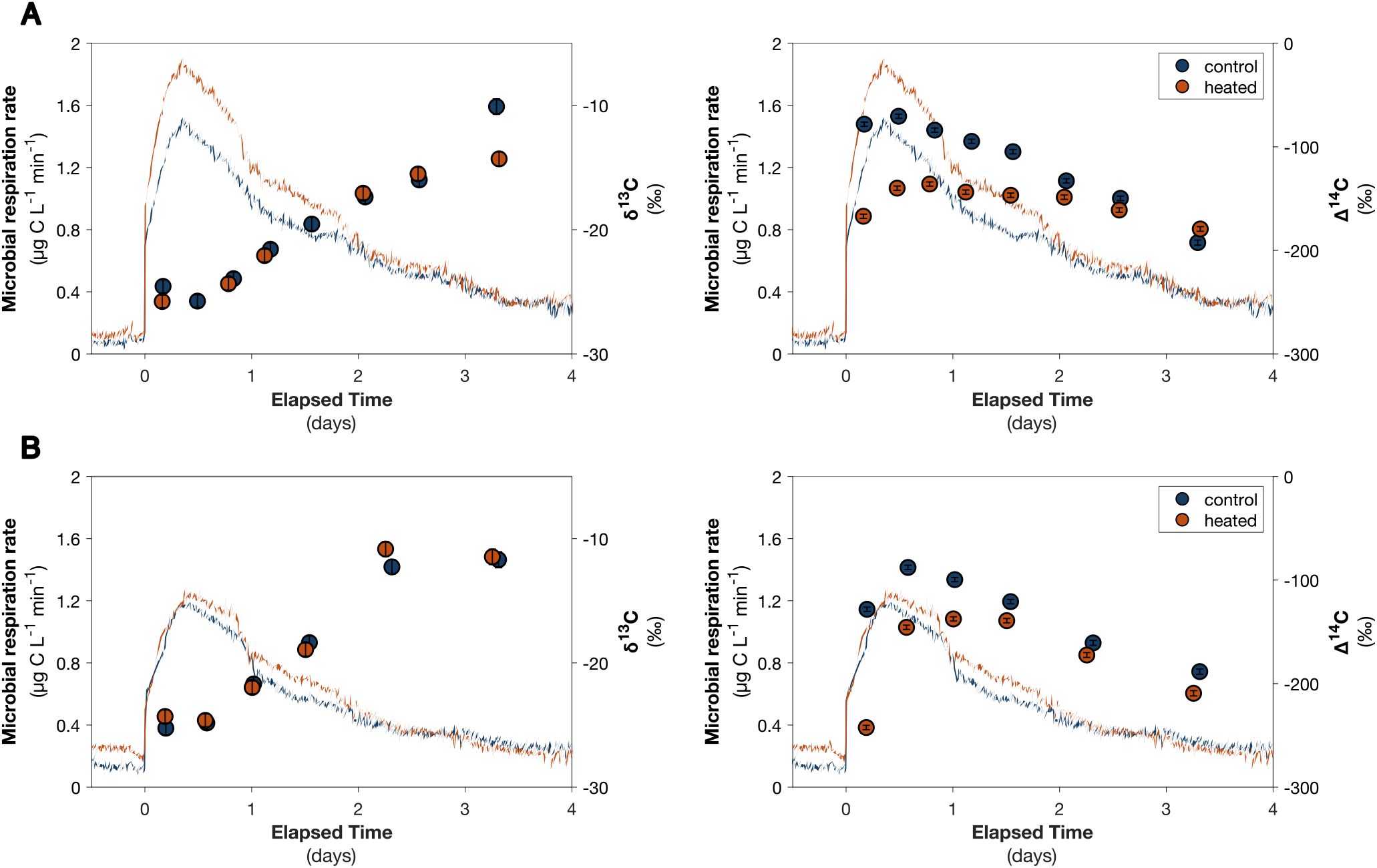
Microbial CO_2_ respiration rates measured during incubations of *Pseudoalteromonas* sp. 3D05 with (A) shallow and (B) deep Guaymas Basin sediment collected from site U1547. Blue curves represent incubations with control sediments, whereas orange curves represent incubations with heated sediments. Corresponding δ^13^C and Δ^14^C concentrations of respired CO_2_ are shown for control (blue circles) and heated (orange circles) incubations.

### 3.3 Isotopic signatures of microbially respired CO_2_

A total of 6 to 8 successive fractions of respired CO_2_ were collected during each incubation for natural abundance C isotopic analysis (δ^13^C and Δ^14^C) to assess the source and age of OM contributing to microbial respiration (Table S5). There was no significant difference observed in the δ^13^C values of microbially respired CO_2_ between control and heated sediment incubations (paired t-test; Table S5). Across all incubations, δ^13^C values showed a progressive enrichment over time, ranging from −28 ± 0.5‰ to −10 ± 0.6‰ (Figures 4, S2). In contrast, Δ^14^C values of respired CO_2_ were different between control and heated sediment incubations, with heating leading to consistently more ^14^C-depleted CO_2_ respired during the first 24 h of incubation. On average, respired CO_2_ was ≥40‰ more depleted in ^14^C during this initial period, indicating that heating altered the radiocarbon age of the OM initially fueling respiration. For sediments from site U1547, Δ^14^C values after the first 24 h ranged from −71 to −83‰ (±2‰) in shallow control incubations and from −136 to −167‰ in shallow heated incubations. In deep U1547 sediments, Δ^14^C values ranged from −88 to −128‰ in control incubations and from −146 to −242‰ following heating (Figure 4; Table S5). A similar pattern was observed in sediments from site U1545. In shallow U1545 sediments, Δ^14^C values of respired CO_2_ during the first 24 h ranged from −33 to −36‰ in control incubations and from −85 to −90‰ in heated incubations. In deep U1545 sediments, control incubations yielded Δ^14^C values of −61 to −73‰, whereas heated incubations ranged from −101 to −138‰ (Figure S2; Table S5).

Since the δ^13^C values of respired CO_2_ were indistinguishable between control and heated sediment incubations, mobilized organic material likely originates from the same broader pool of carbon that is accessed for respiration prior to heating. After the first day, differences in Δ^14^C between control and heated sediment incubations diminished, with the values largely converging by day 2 of incubation. This transient offset suggests that heating renders a fraction of older, more ^14^C depleted material more accessible, which is then preferentially respired early in the incubation. Moreover, the similarity in cumulative CO_2_ yields between control and heated sediment incubations suggests that this pool of ^14^C depleted material is small relative to the total OM pool, such that the lability of the overall OM pool remains unchanged.

## 4. Discussion

A substantial fraction of global marine sediments are exposed to heating due to burial and/or hydrothermal activity. Under these conditions, temperature becomes a major control on the biological processes that govern organic matter transformation in subsurface sediments. Elevated temperatures can promote the generation of low-MW compounds, alter organic-mineral interactions, and potentially modify the composition and bioavailability of the bulk OM pool. Previous studies have shown that heating of marine sediments can generate acetate and other labile organic compounds, but it has remained unclear whether these transformations fundamentally alter the bioavailability of the OM that persists after heating. Understanding how thermal alteration reshapes the balance between labile and more recalcitrant OM pools is critical for understanding constraints on microbial carbon cycling in deeply buried marine sediments.

### 4.1 Sediment freshness governs the capacity for abiotic acetate generation

Our results suggest that the environmental context of sediment (i.e., burial time and hydrothermal history) is likely a key control on the abiotic generation of acetate and supply of substrates to microbial communities. The concentration of acetate in marine sediment porewaters is a function of the balance between biotic and abiotic production and consumption (Burdige and Gardner, 1998). Because these processes occur simultaneously, the controls on abiotic acetate generation are difficult to isolate in natural systems. Our experiments isolated the abiotic component and demonstrated that short-term, low-temperature heating (70 °C for 7 days) leads to the generation of acetate, but that the amount of acetate produced depended strongly on sediment depth and thermal history (Figure 2).

Across all sediment samples, the least amount of acetate was generated during heating of deep sediment from site U1547, which experienced the highest *in situ* temperatures (Teske et al., 2021) prior to laboratory-based heating. This suggests that a history of thermal and diagenetic processing reduces the capacity of sediments to generate acetate through thermal degradation of organic matter. This interpretation is consistent with the measured concentrations of water-extractable DOC in Guaymas Basin sediment porewaters that were lowest in the most extensively heated sediments (Lin et al., 2017; Knoke et al., 2025). We also found acetate generation in marine sediments to be dependent on depth. Previous work by Wellsbury et al. (1997) demonstrated that less acetate was generated from deep sediments incubated at temperatures up to 100°C when compared to incubations with shallow sediments. Taken together, these findings suggest that the pool of OM from which acetate can be generated abiotically is finite. As sediments become progressively altered through burial and diagenetic processing, the capacity to generate acetate, and likely other low-MW organic compounds, is diminished.

### 4.2 Heating alters the age but not the source of respired OM

While the precise mechanisms or pathways are unknown, low-temperature heating is thought to alter OM in two ways. First, it promotes the production of low-MW organic compounds that enter the dissolved phase and can be subsequently transported upward through the sediment column (Lin et al., 2017; Gan et al., 2025). Second, heating at ≥55°C can disrupt organic-mineral associations, releasing more complex protein-like and humic-like material from mineral surfaces (Gan et al., 2025). FT-ICR-MS analysis further shows a loss of organic compounds with 1 or 2 nitrogen atoms during heating to 85°C, likely reflecting cleavage of C-N bonds within humic- or protein-like material (Gan et al., 2025).

Natural abundance carbon isotopic measurements of respired CO_2_ were used to assess how heating altered the OM remineralized by *Pseudoalteromonas* sp. 3D05. When OM is respired by heterotrophic microorganisms, the respired CO_2_ retains the same Δ^14^C and nearly identical δ^13^C signatures as the source material (Hayes, 2001). Consequently, the isotopic composition of respired CO_2_ provides information about both the age and source of the OM being utilized (e.g. Mahmoudi et al., 2020; McNichol et al., 2024; Pearson et al., 2008). Across all sediment samples, δ^13^C values of respired CO_2_ were indistinguishable between control and heated incubations (Figs. 4, S2), indicating that heating did not substantially alter the dominant source of OM fueling microbial respiration. During the first 24 hours of incubation, respired CO_2_ from heated sediments was significantly more depleted in ^14^C than in control incubations (Figures 4, S2). This suggests that short-term, low-temperature heating made an older fraction of OM more accessible. Because the OM pool was not replenished during the bioreactor experiments, substrates consumed early in the incubation are presumed to be the most labile, whereas later CO_2_ fractions reflect respiration of progressively less bioavailable substrates that are only consumed when more labile compounds have been exhausted (Mahmoudi et al., 2017). Thus, the older material liberated during artificial heating, likely through desorption from minerals, is presumed to be more bioavailable (i.e., smaller molecules), since it was consumed within the first 24 hours.

One possible explanation is that OM sorbed to mineral surfaces becomes progressively altered during burial and exposure to heating. A model of organic-mineral interaction in sediments proposed by Kleber et al. (2007) describes a multi-layered structure with two distinct zones of OM interaction (Figure S5). These zones consist of OM sorbed directly to the mineral surface, on top of which lies a second zone of OM electrostatically sorbed to mineral-bound OM through hydrophobic interactions. This creates an “onion-like” coating of OM layers on mineral surfaces (Kleber et al., 2007; Underwood et al., 2024). In this framework, OM directly bound to the mineral surface is more strongly sorbed, while electrostatically bound OM in outer layers is subject to exchange with molecules in the surrounding porewaters. Moreover, the OM bound directly to the mineral surface would likely be older while the electrostatically bound outer layers would consist of younger material.

In our study, sediments experienced moderate (< 55°C) *in situ* heating prior to collection, likely leading to the desorption of material from the younger, outer layers without releasing OM directly sorbed to the mineral surface. Over time, exposure to elevated temperatures *in situ* would deplete the pool of electrostatically sorbed OM. This would leave behind a pool of mineral-sorbed OM with an older average radiocarbon age. When these sediments were artificially heated to 70°C in our experiments, desorption of this inner, mineral-bound OM would have likely made it available to the microorganisms. Moreover, it would be expected that sediments which experienced greater *in situ* heating would be expected to undergo more extensive loss of electrostatically bound, younger outer layers thereby resulting in a larger variation in the Δ^14^C of respired CO_2_ before and after heating. This framework is consistent with our isotopic observations, where the largest Δ^14^C offsets occurred in sediments exposed to the highest *in situ* temperatures. For example, during incubations with deep U1547 sediment (41°C), the difference in Δ^14^C of the first fraction of respired CO_2_ between heated and control incubations was −114‰. With deep U1545 sediment (17°C), however, the difference in Δ^14^C between heated and control incubations was only −64‰ in the first fraction respired (Table S5). This highlights the role of temperature in organic-mineral interactions in marine sediments, as more extensive heating can make a larger fraction of OM sorbed directly to minerals bioavailable.

### 4.3 Short-term heating does not change the overall bioavailability of OM

In our experiments, microbial respiration was primarily supported by the bulk OM pool remaining after generation and removal of volatile compounds such as acetate. Prior to inoculation with *Pseudoalteromonas* sp. 3D05, sediment slurries were sparged for ~ 48h with nitrogen gas. During this process, most of the acetate, and likely other volatile compounds generated from heating, were removed from the bioreactor. Consistent with this, initial acetate concentrations differed little between control and heated incubations (Table S2), despite large differences in acetate production during the separate heating experiments. The bioreactor incubations therefore assessed the bioavailability of the remaining OM pool, including any newly desorbed OM with higher molecular weight retained after sparging.

Despite differences in the Δ^14^C signatures of respired CO_2_, there was no observable difference in the total yield of CO_2_ respired during control and heated sediment incubations (Figure 3). The similarities in respiration rates between control and heated incubations suggest that short-term, low-temperature heating did not measurably alter the bioavailability of the bulk OM pool. However, the seven-day heating experiments conducted here likely capture only the earliest stages of thermal alteration. Marine sediments in hydrothermal environments can experience elevated temperatures for thousands to hundreds of thousands of years. On these geological timescales, repeated thermal alteration and removal of low–MW compounds could progressively deplete the most labile components of sedimentary OM, leaving behind a pool that is increasingly less bioavailable. At the same time, upward mobilization of thermally generated compounds in the sediment column may enhance microbial activity in overlying sediments, even while the availability of labile substrates in deeper horizons is reduced.

## 5. Conclusions

Our study demonstrates that the effects of heating on sedimentary OM depend strongly on sediment depth and prior thermal history. Both shallow and deeply buried marine sediments have the capacity to generate acetate upon low-temperature heating over a short period of time. Although heating generates a small pool of labile compounds, it does not substantially alter the overall bioavailability of the remaining OM, however, it does make a pool of older material from sediments more accessible to subsurface microorganisms. Taken together, our results highlight how temperature can exert complex and variable effects on the bioavailability of OM in marine sediments. Our findings highlight the role of hydrothermal alteration on carbon cycling in marine sediments, through transformation of the organic matter pool and its availability to microorganisms. Understanding these processes is essential for predicting how thermal alteration influences carbon cycling, microbial activity, and the long-term fate of OM in the marine subsurface.

## Acknowledgements

This work was supported by a grant from the U.S. National Science Foundation (award no. 2023656) to SRS and NM and by a Doctoral Research Scholarship from the Fonds de recherche du Québec (#330067) to SMM. This research used samples and data provided by International Ocean Discovery Project (IODP) Expedition 385. We thank the scientists, crew, and technical staff aboard the *JOIDES Resolution* for their dedication and efforts in making this work possible. We are grateful to Dr. Nathan Dalleska for assistance with acetate measurements at Caltech, Dr. Thi Hao Bui for analytical assistance at McGill, and the members of NOSAMS and OSSIC for isotope measurements.

## CRediT authorship contribution statement: SMM

Writing – review & editing, Writing – original draft, Visualization, Investigation, Formal analysis, Data curation, Conceptualization. **SSW:** Writing – review & editing, Funding acquisition, Conceptualization. **AT:** Writing – review & editing. **NM:** Writing – review & editing, Writing – original draft, Supervision, Methodology, Investigation, Funding acquisition, Conceptualization.

## Appendix A. Supplemental Material

The supplementary material includes 4 Figures and 6 Tables. Figures present (1) a simplified schematic of experiments carried out with sediment samples, (2) microbial CO_2_ respiration rates and corresponding δ^13^C and Δ^14^C concentrations measured during incubations with U1545 sediment, (3) microbial CO_2_ respiration rates and cell densities measured during incubations U1547 sediment, (4) microbial CO_2_ respiration rates and cell densities measured during incubations with U1545 sediment, and (5) schematic diagram of OM-mineral interactions. Tables provide (1) sediment sample depth and *in situ* temperature, (2) mass of sediment incubated and initial bioreactor acetate concentration for all incubations, (3) percent total organic carbon (TOC), mass of sediment incubated, mass of carbon (C) respired, and percent TOC respired during all incubations, (4) cell densities measured during incubations, (5) δ^13^C and Δ^14^C concentrations of CO_2_ respired during incubations, and (6) p-values for paired t-tests comparing δ^13^C values from respired CO_2_ between control and heated sediment incubations.

