## Supplemental Information for "The effects of thermal alteration on organic matter bioavailability in deeply buried marine sediments"

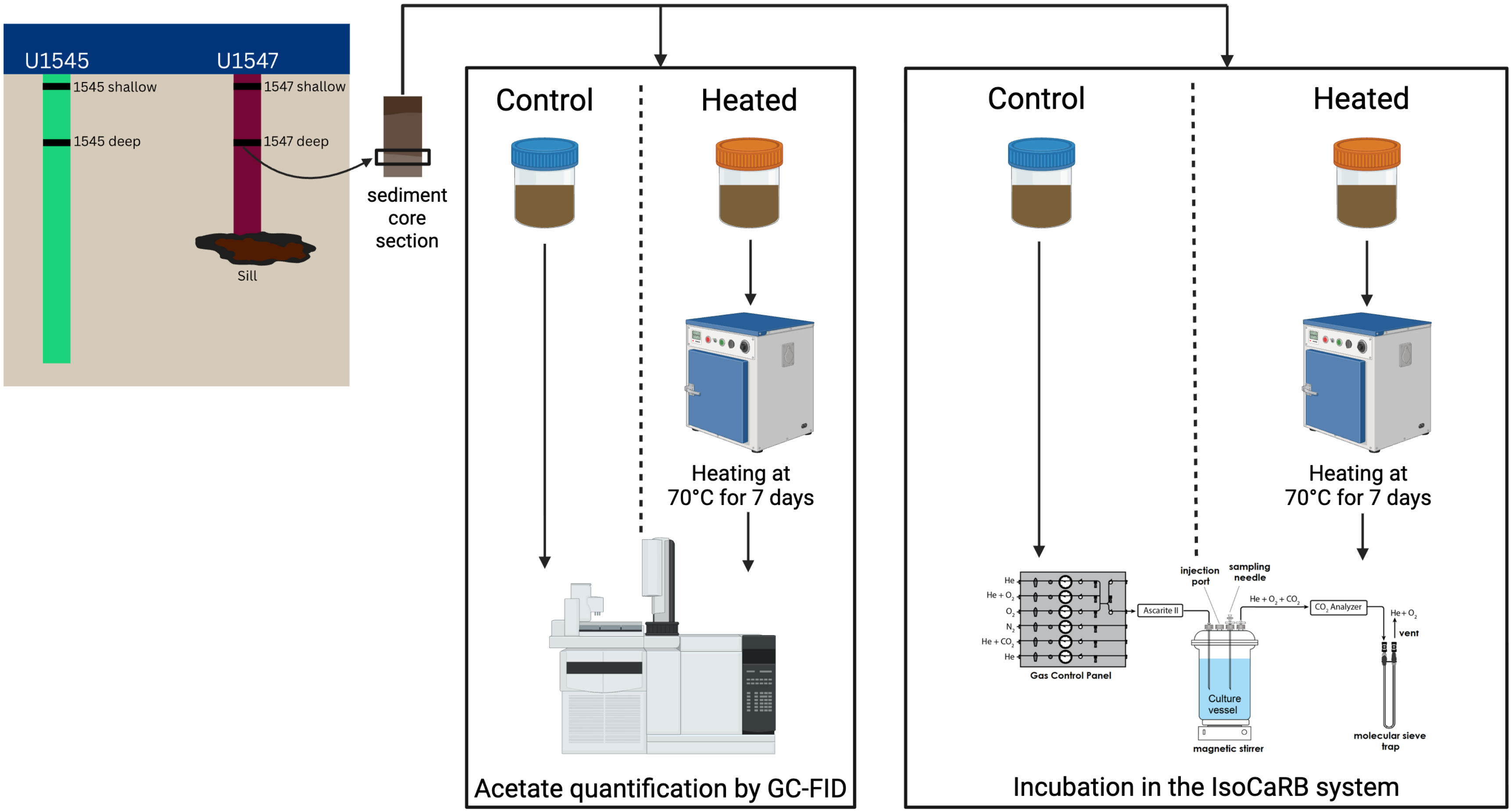


**Figure S1.** Simplified schematic of experiments carried out for each sediment sample. Both heated and control sediment aliquots were taken from the same, homogenized sediment core section. All sediment used in this study was sterilized and freeze dried before use in experiments. Heated and control sediment slurries were filtered and analyzed by GC-FID to quantify acetate concentrations. Two separate aliquots of the same sediment core section were used for incubation in the IsoCaRB system. One aliquot was heated, the other left untreated as a control. Each aliquot was incubated separately in the IsoCaRB with *Pseudoalteromonas* sp. 3D05. This image was created using BioRender.com.

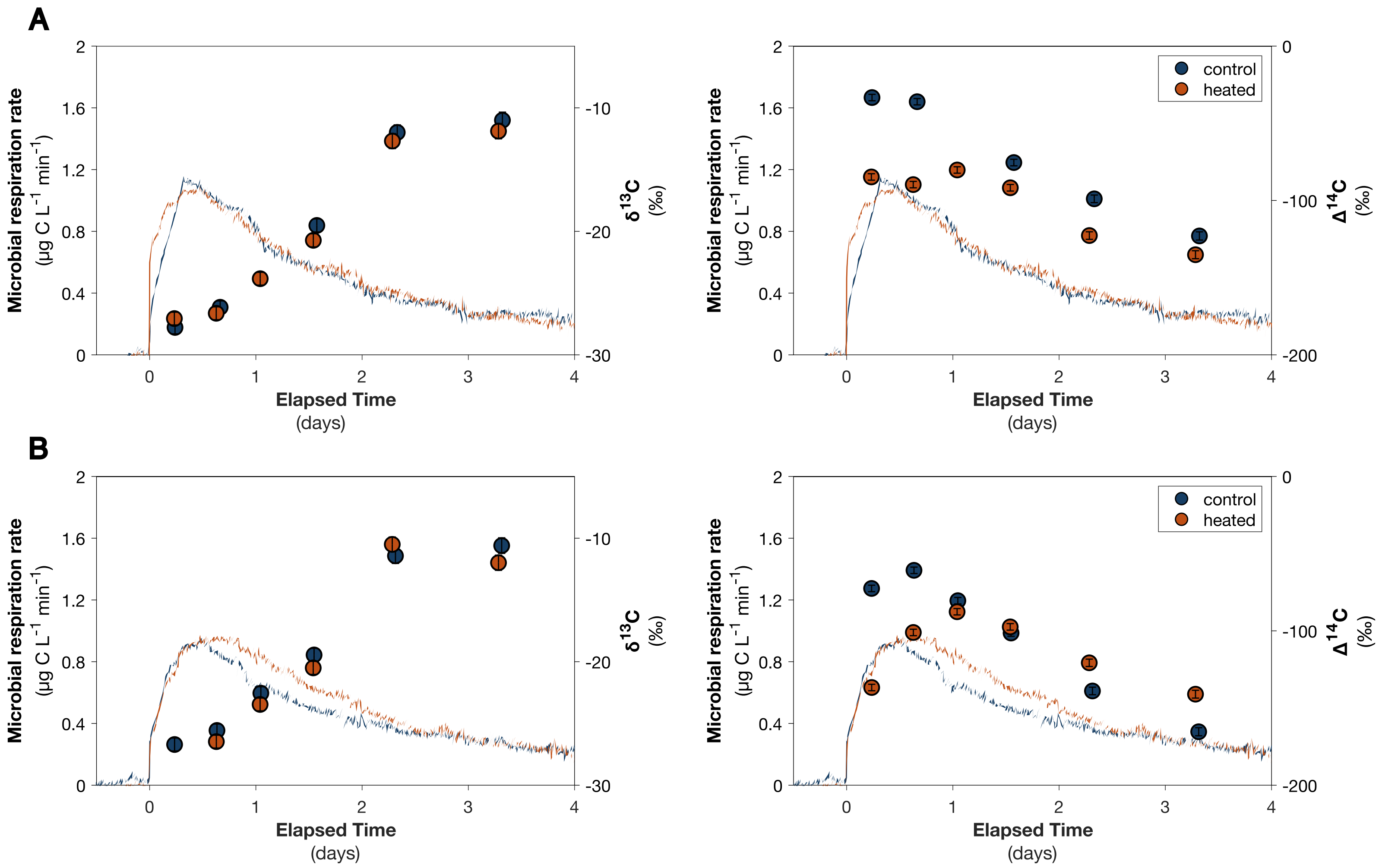


**Figure S2.** Microbial $CO_2$ respiration rates measured during incubations of *Pseudoalteromonas* sp. 3D05 with (A) shallow and (B) deep Guaymas Basin sediment collected from site U1545. Blue curves represent incubations with control sediments, whereas orange curves represent incubations with heated sediments. Corresponding $\delta^{13}C$ and $\Delta^{14}C$ concentrations of respired $CO_2$ are shown for control (blue circles) and heated (orange circles) incubations.

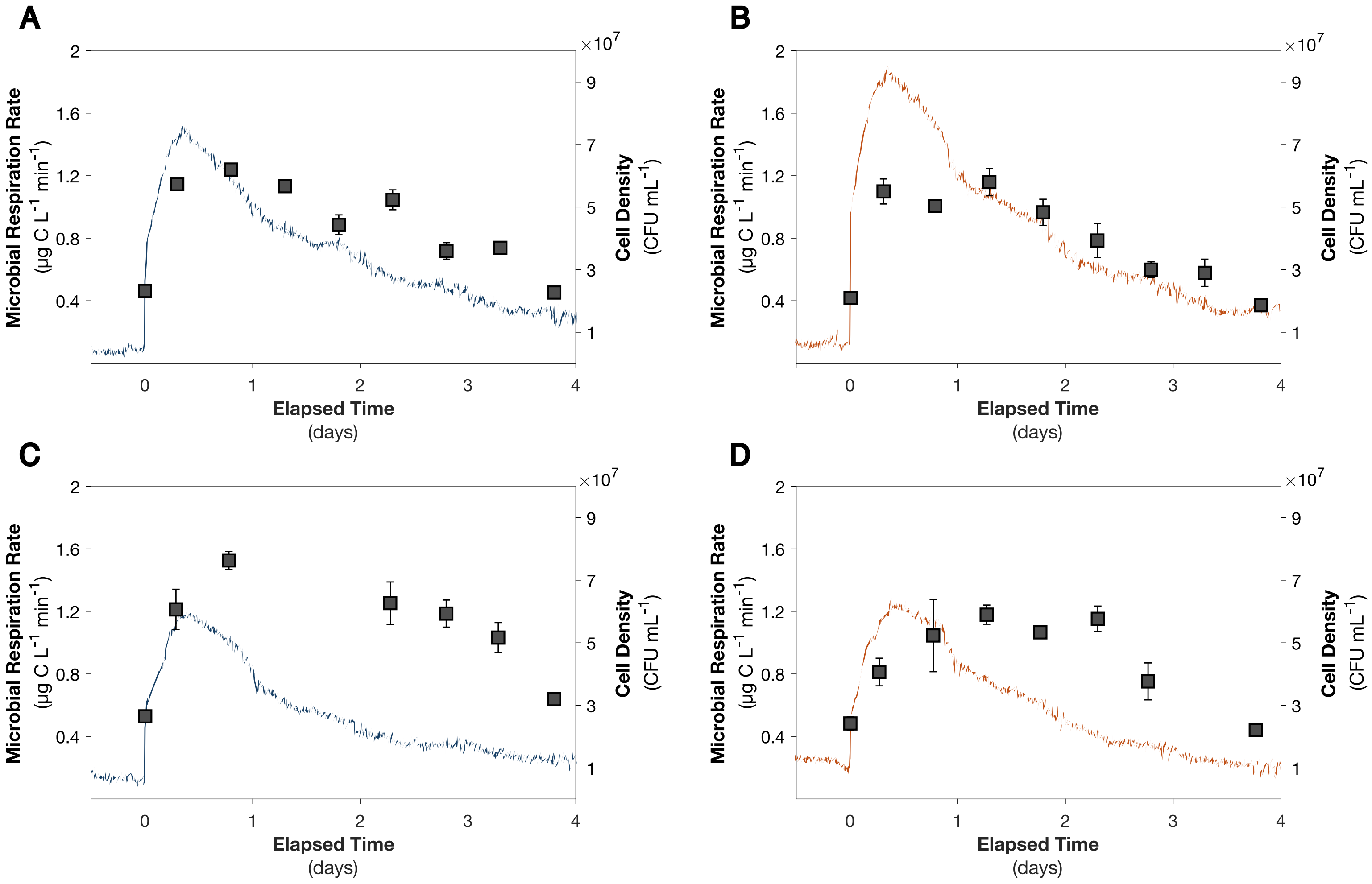


**Figure S3.** Microbial respiration rates (blue and orange lines) and cell densities (gray squares) measured during incubations of *Pseudoalteromonas* sp. 3D05 with (A) shallow, control, site U1547 sediment; (B) shallow, heated, site U1547 sediment; (C) deep, control, site U1547 sediment; (D) deep, heated, site U1547 sediment. Error bars represent the standard deviation between CFU/ml measurements from triplicate plates.

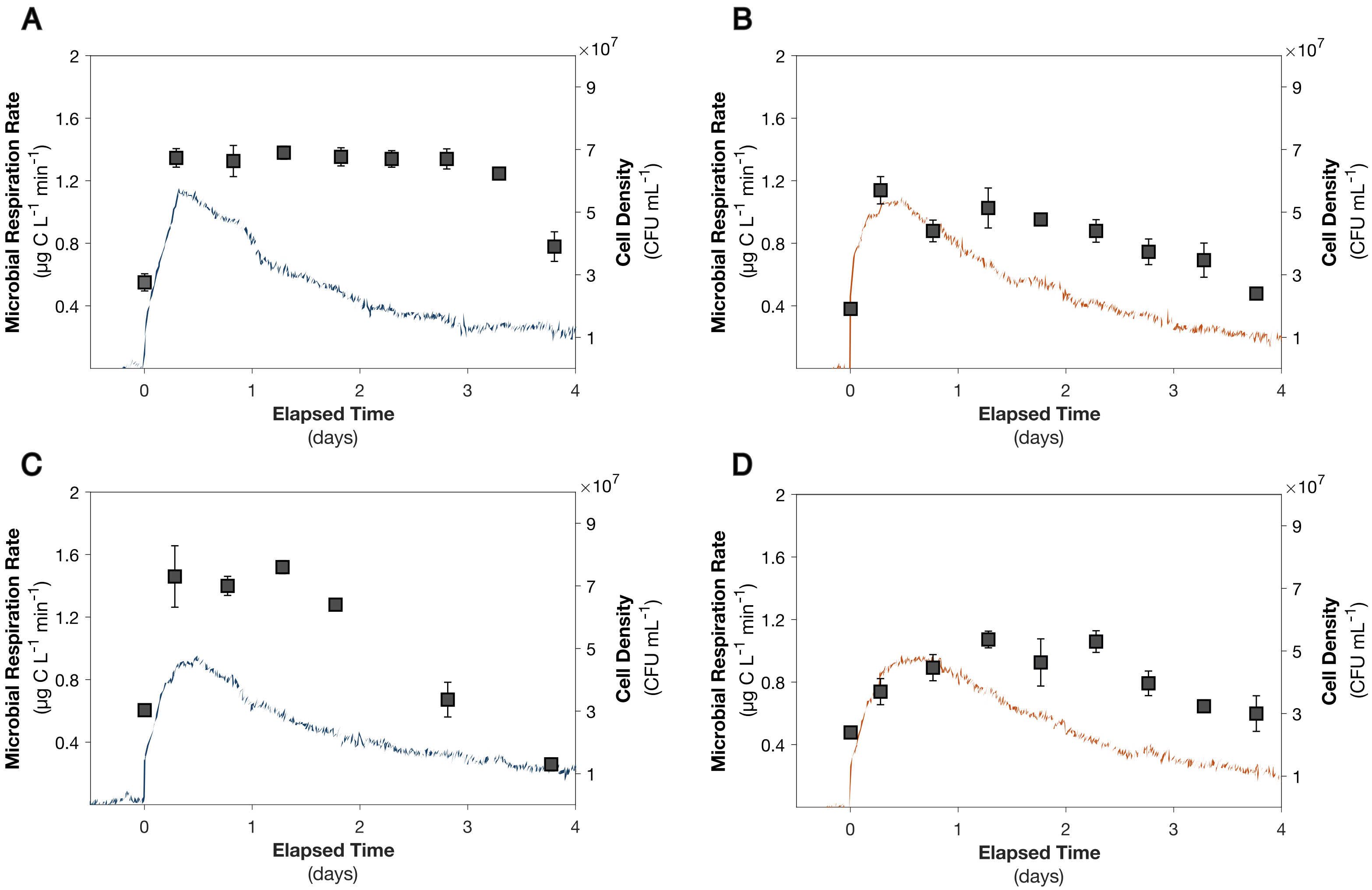


**Figure S4.** Microbial respiration rates (blue and orange lines) and cell densities (gray squares) measured during incubations of *Pseudoalteromonas* sp. 3D05 with (A) shallow, control, site U1545 sediment; (B) shallow, heated, site U1545 sediment; (C) deep, control, site U1545 sediment; (D) deep, heated, site U1545 sediment. Error bars represent the standard deviation between CFU/ml measurements from triplicate plates.

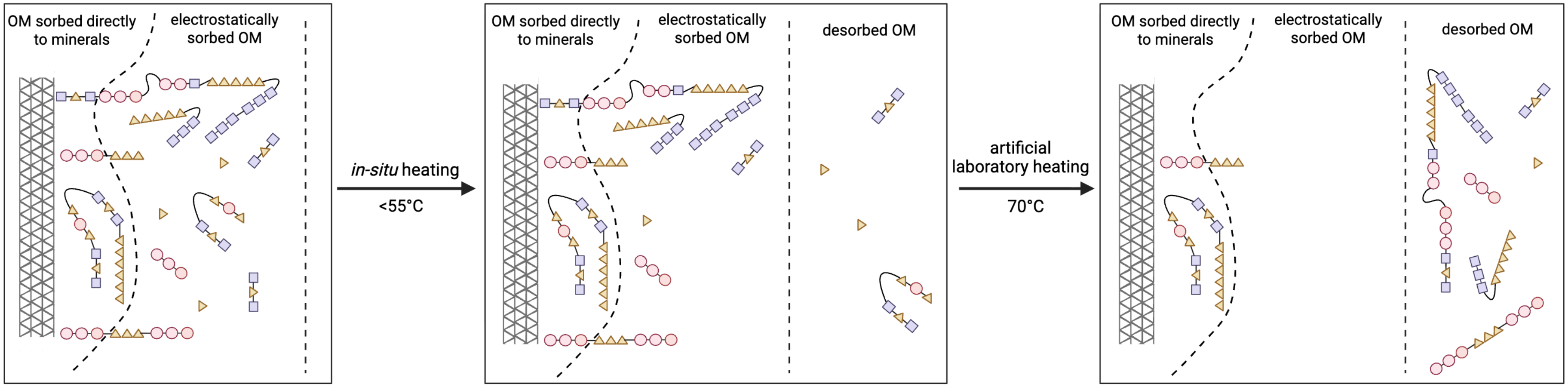


**Figure S5.** Schematic diagram representing the zones of organic-mineral interactions, adapted from Kleber et al., 2007. In the zone closest to the mineral surface (gray cross-hatched rectangle) molecules are sorbed directly to the mineral surface. Just beyond this zone, molecules are electrostatically sorbed to mineral-bound OM, held mainly by hydrophobic interactions. After *in situ* heating (<55°C), molecules sorbed electrostatically can be released to the surrounding porewaters, while molecules sorbed directly to the mineral surface remain bound. After heating at higher temperatures in the laboratory (70°C), both electrostatically sorbed molecules and molecules bound directly to the mineral surface can be desorbed and released to the surrounding porewaters, becoming more available for microbial consumption or transport upward in the sediment column. This image was created using BioRender.com.

### SUPPLEMENTAL TABLES

**Table S1.** Depth and *in situ* temperature of shallow and deep sediment core sections from sites U1547 and U1545.

| Site | Sample | Depth (mbsf) | Temperature (°C) |
|---|---|---|---|
| U1547 | shallow | 8.9 | 13 |
| | deep | 56 | 41 |
| U1545 | shallow | 4.3 | 7 |
| | deep | 57 | 17 |

**Table S2.** Mass of sediment incubated and initial acetate concentration measured from the bioreactor slurry during control and heated sediment incubations. Acetate concentrations were determined by GC-FID from a subsample of the bioreactor slurry.

| | Sediment Mass Incubated (g) | Initial Acetate Concentration (μM) |
|---|---|---|
| **U1547 shallow** | | |
| Control | 45 | 56 ± 1 |
| Heated | 45 | 40 ± 5 |
| **U1547 deep** | | |
| Control | 35 | 21 ± 3 |
| Heated | 35 | 45 ± 2 |
| **U1545 shallow** | | |
| Control | 22 | 43 ± 4 |
| Heated | 22 | 28 ± 5 |
| **U1545 deep** | | |
| Control | 28 | 16 ± 1 |
| Heated | 28 | 32 ± 3 |

**Table S3.** Percent total organic carbon (TOC), mass of sediment incubated, mass of carbon (C) respired, and percent TOC respired during control and heated sediment incubations with *Pseudoalteromonas* sp. 3D05.

| | TOC (%) | Sediment Mass Incubated (g) | C Respired (mg) | % TOC Respired |
|---|---|---|---|---|
| **U1547 shallow** | | | | |
| Control | 3.52 | 45 | 8.7 ± 0.01 | 0.55 ± 0.01 |
| Heated | 3.52 | 45 | 10.2 ± 0.01 | 0.64 ± 0.01 |
| **U1547 deep** | | | | |
| Control | 2.61 | 35 | 6.7 ± 0.01 | 0.73 ± 0.03 |
| Heated | 2.61 | 35 | 6.9 ± 0.01 | 0.76 ± 0.03 |
| **U1545 shallow** | | | | |
| Control | 2.26 | 22 | 6.2 ± 0.01 | 1.25 ± 0.01 |
| Heated | 2.26 | 22 | 6.3 ± 0.01 | 1.26 ± 0.01 |
| **U1545 deep** | | | | |
| Control | 1.77 | 28 | 5.7 ± 0.01 | 1.14 ± 0.05 |
| Heated | 1.77 | 28 | 6.2 ± 0.01 | 1.24 ± 0.05 |

**Table S4.** Cell densities of *Pseudoalteromonas* sp. 3D05 measured during incubation experiments in the IsoCaRB system. Cell density was measured by manual counting of colony forming units (CFUs) on agar plates from subsamples of the sediment slurry.

| | Elapsed Time (days) | *Pseudoalteromonas* sp. 3D05 (CFU/mL) |
|---|---|---|
| **U1547 shallow control** | | |
| | 0.0 | $2.3 \pm 0.05 \times 10^7$ |
| | 0.3 | $5.7 \pm 0.1 \times 10^7$ |
| | 0.8 | $6.2 \pm 0.2 \times 10^7$ |
| | 1.3 | $5.7 \pm 0.2 \times 10^7$ |
| | 1.8 | $4.4 \pm 0.3 \times 10^7$ |
| | 2.3 | $5.2 \pm 0.3 \times 10^7$ |
| | 2.8 | $3.6 \pm 0.3 \times 10^7$ |
| | 3.3 | $3.7 \pm 0.2 \times 10^7$ |
| | 3.8 | $2.3 \pm 0.006 \times 10^7$ |
| | 4.3 | $1.4 \pm 0.2 \times 10^7$ |
| | 4.8 | $4.6 \pm 0.6 \times 10^6$ |
| | 5.3 | $5.8 \pm 0.5 \times 10^6$ |
| | 5.8 | $2.6 \pm 0.3 \times 10^6$ |
| **U1547 shallow heated** | | |
| | 0.0 | $2.1 \pm 0.06 \times 10^7$ |
| | 0.3 | $5.5 \pm 0.4 \times 10^7$ |
| | 0.8 | $5.0 \pm 0.09 \times 10^7$ |
| | 1.3 | $5.8 \pm 0.4 \times 10^7$ |
| | 1.8 | $4.8 \pm 0.4 \times 10^7$ |
| | 2.3 | $3.9 \pm 0.6 \times 10^7$ |
| | 2.8 | $3.0 \pm 0.3 \times 10^7$ |
| | 3.3 | $2.9 \pm 0.4 \times 10^7$ |
| | 3.8 | $1.9 \pm 0.09 \times 10^7$ |
| | 4.3 | $1.5 \pm 0.06 \times 10^7$ |
| | 4.8 | $5.7 \pm 0.6 \times 10^6$ |
| | 5.3 | $6.4 \pm 0.7 \times 10^6$ |
| | 5.8 | $2.0 \pm 0.3 \times 10^6$ |
| **U1547 deep control** | | |
| | 0.0 | $2.7 \pm 0.09 \times 10^7$ |
| | 0.3 | $6.1 \pm 0.6 \times 10^7$ |
| | 0.8 | $7.6 \pm 0.3 \times 10^7$ |
| | 1.3 | |
| | 1.8 | |
| | 2.3 | $6.3 \pm 0.7 \times 10^7$ |
| | 2.8 | $5.9 \pm 0.4 \times 10^7$ |
| | 3.3 | $5.2 \pm 0.5 \times 10^7$ |
| | 3.8 | $3.2 \pm 0.1 \times 10^7$ |
| | 4.3 | $3.0 \pm 0.06 \times 10^7$ |
| | 4.8 | $2.1 \pm 0.06 \times 10^7$ |
| | 5.3 | $1.7 \pm 0.01 \times 10^7$ |

| | | |
|---|---|---|
| | 5.8 | $6.2 \pm 0.7 \times 10^6$ |
| **U1547 deep heated** | | |
| | 0.0 | $2.4 \pm 0.2 \times 10^7$ |
| | 0.3 | $4.1 \pm 0.4 \times 10^7$ |
| | 0.8 | $5.2 \pm 1 \times 10^7$ |
| | 1.3 | $5.9 \pm 0.3 \times 10^7$ |
| | 1.8 | $5.3 \pm 0.2 \times 10^7$ |
| | 2.3 | $5.8 \pm 0.4 \times 10^7$ |
| | 2.8 | $3.8 \pm 0.6 \times 10^7$ |
| | 3.3 | |
| | 3.8 | $2.2 \pm 0.1 \times 10^7$ |
| | 4.3 | $2.5 \pm 0.1 \times 10^7$ |
| | 4.8 | $1.3 \pm 0.03 \times 10^7$ |
| | 5.3 | $1.7 \pm 0.1 \times 10^7$ |
| | 5.8 | $9.0 \pm 0.6 \times 10^6$ |
| **U1545 shallow control** | | |
| | 0.0 | $2.8 \pm 0.3 \times 10^7$ |
| | 0.3 | $6.7 \pm 0.3 \times 10^7$ |
| | 0.8 | $6.6 \pm 0.5 \times 10^7$ |
| | 1.3 | $6.9 \pm 0.2 \times 10^7$ |
| | 1.8 | $6.8 \pm 0.3 \times 10^7$ |
| | 2.3 | $6.7 \pm 0.3 \times 10^7$ |
| | 2.8 | $6.7 \pm 0.3 \times 10^7$ |
| | 3.3 | $6.2 \pm 0.2 \times 10^7$ |
| | 3.8 | $3.9 \pm 0.5 \times 10^7$ |
| | 4.3 | $4.5 \pm 0.7 \times 10^7$ |
| | 4.8 | $1.7 \pm 0.3 \times 10^7$ |
| | 5.3 | $2.9 \pm 0.02 \times 10^7$ |
| | 5.8 | $5.6 \pm 0.8 \times 10^6$ |
| **U1545 shallow heated** | | |
| | 0.0 | $1.9 \pm 0.1 \times 10^7$ |
| | 0.3 | $5.7 \pm 0.4 \times 10^7$ |
| | 0.8 | $4.4 \pm 0.4 \times 10^7$ |
| | 1.3 | $5.1 \pm 0.6 \times 10^7$ |
| | 1.8 | $4.8 \pm 0.07 \times 10^7$ |
| | 2.3 | $4.4 \pm 0.4 \times 10^7$ |
| | 2.8 | $3.7 \pm 0.4 \times 10^7$ |
| | 3.3 | $3.5 \pm 0.6 \times 10^7$ |
| | 3.8 | $2.4 \pm 0.2 \times 10^7$ |
| | 4.3 | $2.2 \pm 0.3 \times 10^7$ |
| | 4.8 | $8.8 \pm 0.5 \times 10^6$ |
| | 5.3 | $7.7 \pm 0.8 \times 10^6$ |
| | 5.8 | $3.2 \pm 0.3 \times 10^6$ |

| | | |
|---|---|---|
| **U1545 deep control** | | |
| | 0.0 | $3.0 \pm 0.1 \times 10^7$ |
| | 0.3 | $7.3 \pm 1 \times 10^7$ |
| | 0.8 | $7.0 \pm 0.3 \times 10^7$ |
| | 1.3 | $7.6 \pm 0.2 \times 10^7$ |
| | 1.8 | $6.4 \pm 0.2 \times 10^7$ |
| | 2.3 | |
| | 2.8 | $3.4 \pm 0.6 \times 10^7$ |
| | 3.3 | |
| | 3.8 | $1.3 \pm 0.2 \times 10^7$ |
| | 4.3 | $7.3 \pm 0.1 \times 10^6$ |
| | 4.8 | $7.8 \pm 0.9 \times 10^6$ |
| | 5.3 | $1.0 \pm 0.1 \times 10^7$ |
| | 5.8 | $3.4 \pm 0.7 \times 10^6$ |
| **U1545 deep heated** | | |
| | 0.0 | $2.4 \pm 0.2 \times 10^7$ |
| | 0.3 | $3.7 \pm 0.4 \times 10^7$ |
| | 0.8 | $4.5 \pm 0.4 \times 10^7$ |
| | 1.3 | $5.4 \pm 0.3 \times 10^7$ |
| | 1.8 | $4.6 \pm 0.8 \times 10^7$ |
| | 2.3 | $5.3 \pm 0.4 \times 10^7$ |
| | 2.8 | $4.0 \pm 0.4 \times 10^7$ |
| | 3.3 | $3.2 \pm 0.2 \times 10^7$ |
| | 3.8 | $3.0 \pm 0.6 \times 10^7$ |
| | 4.3 | $2.0 \pm 0.6 \times 10^7$ |
| | 4.8 | $9.5 \pm 0.6 \times 10^6$ |
| | 5.3 | $9.3 \pm 1 \times 10^6$ |
| | 5.8 | $7.7 \pm 1 \times 10^6$ |

**Table S5.** Masses and isotopic values of microbially-respired $CO_2$ collected during control and heated incubations of *Pseudoalteromoas* sp. 3D05 with shallow and deep Guaymas Basin sediment.

| | Fraction # | Collection Times (days) | Collection Duration (hr) | Mass $CO_2$ (µg C) | $\delta^{13}C$ (‰) | $\Delta^{14}C$ (‰) |
|---|---|---|---|---|---|---|
| **Control Sediment** | | | | | | |
| *1545B 2H 4* | 1 | 0.0-0.5 | 11.5 | 1121 ± 3 | -28 ± 0.5 | -33 ± 2 |
| | 2 | 0.5-0.8 | 8.9 | 1212 ± 3 | -26 ± 0.5 | -36 ± 2 |
| | 3 | 0.8-1.3 | 11.0 | 1150 ± 3 | | |
| | 4 | 1.3-1.8 | 12.8 | 971 ± 2 | -20 ± 0.5 | -75 ± 2 |
| | 5 | 1.8-2.8 | 23.5 | 550 ± 1 | -12 ± 0.6 | -99 ± 2 |
| | 6 | 2.8-3.8 | 24.0 | 431 ± 1 | -11 ± 0.7 | -123 ± 3 |
| | 7 | 3.8-4.8 | 24.0 | 406 ± 1 | | |
| | 8 | 4.8-5.8 | 24.1 | 285 ± 1 | | |
| *1545B 7H 4* | 1 | 0.0-0.5 | 11.3 | 1077 ± 2 | -27 ± 0.5 | -73 ± 2 |
| | 2 | 0.5-0.8 | 7.9 | 912 ± 2 | -26 ± 0.5 | -61 ± 2 |
| | 3 | 0.8-1.3 | 12.0 | 1115 ± 3 | -23 ± 0.5 | -80 ± 2 |
| | 4 | 1.3-1.8 | 12.0 | 862 ± 2 | -19 ± 0.5 | -101 ± 2 |
| | 5 | 1.8-2.8 | 24.8 | 531 ± 1 | -11 ± 0.6 | -139 ± 2 |
| | 6 | 2.8-3.8 | 23.2 | 491 ± 1 | -11 ± 0.6 | -165 ± 3 |
| | 7 | 3.8-4.8 | 24.0 | 407 ± 1 | | |
| | 8 | 4.8-5.8 | 24.0 | 281 ± 1 | | |
| *1547B 1H 3* | 1 | 0.0-0.3 | 8.2 | 1120 ± 3 | -25 ± 0.5 | -78 ± 2 |
| | 2 | 0.3-0.6 | 7.3 | 1290 ± 3 | -26 ± 0.5 | -71 ± 2 |
| | 3 | 0.6-1.0 | 8.9 | 1347 ± 3 | -24 ± 0.5 | -84 ± 2 |
| | 4 | 1.0-1.3 | 7.7 | 949 ± 2 | -22 ± 0.5 | -95 ± 2 |
| | 5 | 1.3-1.8 | 10.8 | 1085 ± 3 | -19 ± 0.5 | -105 ± 2 |
| | 6 | 1.8-2.3 | 13.2 | 969 ± 2 | -17 ± 0.5 | -133 ± 2 |
| | 7 | 2.3-2.8 | 11.2 | 705 ± 2 | -16 ± 0.5 | -150 ± 2 |
| | 8 | 2.8-3.8 | 23.6 | 509 ± 1 | -10 ± 0.6 | -192 ± 2 |
| | 9 | 3.8-4.8 | 24.3 | 342 ± 1 | -8 ± 0.7 | -234 ± 3 |
| | 10 | 4.8-5.8 | 24.4 | 354 ± 1 | -9 ± 0.7 | -285 ± 3 |
| *1547B 7H 3* | 1 | 0.0-0.4 | 9.4 | 1109 ± 3 | -25 ± 0.5 | -128 ± 2 |
| | 2 | 0.4-0.8 | 9.0 | 1351 ± 3 | -25 ± 0.5 | -88 ± 2 |
| | 3 | 0.8-1.3 | 12.0 | 1322 ± 3 | -22 ± 0.5 | -100 ± 2 |
| | 4 | 1.3-1.8 | 13.1 | 1007 ± 2 | -18 ± 0.5 | -121 ± 2 |
| | 5 | 1.8-2.8 | 23.9 | 666 ± 2 | -12 ± 0.6 | -161 ± 2 |
| | 6 | 2.8-3.8 | 24.0 | 463 ± 1 | -12 ± 0.6 | -188 ± 3 |
| | 7 | 3.8-4.8 | 24.0 | 428 ± 1 | -12 ± 0.7 | -220 ± 3 |
| | 8 | 4.8-5.8 | 24.3 | 337 ± 1 | -14 ± 0.7 | -269 ± 3 |

| | | | | | | |
|---|---|---|---|---|---|---|
| **Heated Sediment** | | | | | | |
| *1545B 2H 4* | 1 | 0.0-0.5 | 11.2 | 1341 ± 3 | -27 ± 0.5 | -85 ± 2 |
| | 2 | 0.5-0.8 | 7.7 | 1016 ± 2 | -27 ± 0.5 | -90 ± 2 |
| | 3 | 0.8-1.3 | 12.2 | 1302 ± 3 | -24 ± 0.5 | -80 ± 2 |
| | 4 | 1.3-1.8 | 11.8 | 918 ± 2 | -21 ± 0.5 | -92 ± 2 |
| | 5 | 1.8-2.8 | 24.0 | 559 ± 1 | -13 ± 0.6 | -123 ± 2 |
| | 6 | 2.8-3.8 | 24.0 | 513 ± 1 | -12 ± 0.6 | -135 ± 3 |
| | 7 | 3.8-4.8 | 24.0 | 338 ± 1 | | |
| | 8 | 4.8-5.8 | 24.0 | 273 ± 1 | | |
| *1545B 7H 4* | 1 | 0.0-0.5 | 11.3 | 1035 ± 2 | | -138 ± 2 |
| | 2 | 0.5-0.8 | 7.5 | 909 ± 2 | -26 ± 0.5 | -101 ± 2 |
| | 3 | 0.8-1.3 | 12.3 | 1365 ± 3 | -23 ± 0.5 | -88 ± 2 |
| | 4 | 1.3-1.8 | 11.7 | 1023 ± 2 | -21 ± 0.5 | -97 ± 2 |
| | 5 | 1.8-2.8 | 24.0 | 603 ± 1 | -11 ± 0.6 | -121 ± 2 |
| | 6 | 2.8-3.8 | 24.0 | 547 ± 1 | -12 ± 0.6 | -141 ± 3 |
| | 7 | 3.8-4.8 | 24.0 | 389 ± 1 | | |
| | 8 | 4.8-5.8 | 24.0 | 294 ± 1 | | |
| *1547B 1H 3* | 1 | 0.0-0.3 | 7.8 | 1267 ± 3 | -26 ± 0.5 | -167 ± 2 |
| | 2 | 0.3-0.6 | 7.3 | 1509 ± 3 | | -140 ± 2 |
| | 3 | 0.6-0.9 | 7.3 | 1296 ± 3 | -24 ± 0.5 | -136 ± 2 |
| | 4 | 0.9-1.3 | 8.9 | 1284 ± 3 | -22 ± 0.5 | -144 ± 2 |
| | 5 | 1.3-1.8 | 11.4 | 1263 ± 3 | | -147 ± 2 |
| | 6 | 1.8-2.3 | 12.7 | 1081 ± 2 | -17 ± 0.5 | -149 ± 2 |
| | 7 | 2.3-2.8 | 12.0 | 779 ± 2 | -16 ± 0.5 | -161 ± 2 |
| | 8 | 2.8-3.8 | 24.5 | 866 ± 2 | -14 ± 0.6 | -179 ± 2 |
| | 9 | 3.8-4.8 | 23.6 | 410 ± 1 | -9 ± 0.7 | -212 ± 3 |
| | 10 | 4.8-5.8 | 24.5 | 454 ± 1 | | -245 ± 3 |
| *1547B 7H 3* | 1 | 0.0-0.4 | 9.1 | 1075 ± 2 | -24 ± 0.5 | -242 ± 2 |
| | 2 | 0.4-0.8 | 9.0 | 1388 ± 3 | -25 ± 0.5 | -146 ± 2 |
| | 3 | 0.8-1.3 | 12.1 | 1523 ± 3 | -22 ± 0.5 | -138 ± 2 |
| | 4 | 1.3-1.8 | 11.9 | 1037 ± 2 | -19 ± 0.5 | -139 ± 2 |
| | 5 | 1.8-2.8 | 24.0 | 630 ± 1 | -11 ± 0.6 | -172 ± 2 |
| | 6 | 2.8-3.8 | 24.0 | 533 ± 1 | -11 ± 0.6 | -209 ± 3 |
| | 7 | 3.8-4.8 | 24.0 | 422 ± 1 | | |
| | 8 | 4.8-5.8 | 24.7 | 323 ± 1 | | |

**Table S6.** P-value for paired t-tests comparing $\delta^{13}C$ values from respired $CO_2$ between control and heated sediment incubations.

| Sediment sample | p-value |
|---|---|
| U1547 shallow | 0.2441 |
| U1547 deep | 0.3297 |
| U1545 shallow | 0.1951 |
| U1545 deep | 0.1783 |
